# A chromosome-scale genome assembly of the false clownfish, *Amphiprion ocellaris*

**DOI:** 10.1101/2022.01.16.476524

**Authors:** Taewoo Ryu, Marcela Herrera, Billy Moore, Michael Izumiyama, Erina Kawai, Vincent Laudet, Timothy Ravasi

## Abstract

**Background:** The false clownfish *Amphiprion ocellaris* is a popular fish species and an emerging model organism for studying the ecology, evolution, adaptation, and developmental biology of reef fishes. Despite this, high-quality genomic resources for this species are scarce, hindering advanced genomic analyses. Leveraging the power of PacBio long-read sequencing and Hi-C chromosome conformation capture techniques, we constructed a high-quality chromosome-scale genome assembly for the clownfish *A. ocellaris*.

**Results:** The initial genome assembly comprised of 1,551 contigs of 861.42 Mb, with an N50 of 863.85 kb. Hi-C scaffolding of the genome resulted in 24 chromosomes containing 856.61 Mb. The genome was annotated with 26,797 protein-coding genes and had 96.62 % completeness of conserved actinopterygian genes, making this genome the most complete and high quality among published anemonefish genomes. Transcriptomic analysis identified tissue-specific gene expression patterns, with the brain and optic lobe having the largest number of expressed genes. Further, comparative genomic analysis revealed 91 genome elements conserved only in *A. ocellaris* and its sister species *Amphiprion percula*, and not in other anemonefish species. These elements are close to genes that are involved in various nervous system functions and exhibited distinct expression patterns in brain tissue, potentially highlighting the genetic toolkits involved in lineage-specific divergence and behaviors of the clownfish branch.

**Conclusions:** Overall, our study provides the highest quality *A. ocellaris* genome assembly and annotation to date, whilst also providing a valuable resource for understanding the ecology and evolution of reef fishes.

## DATA DESCRIPTION

### Context

The false clownfish *Amphiprion ocellaris* is one of 28 anemonefishes (from the subfamily Amphiprioninae in the family Pomacentridae) among thousands of tropical marine fish species [1]. Yet, together with its sister species, the orange clownfish *Amphiprion percula*, it is one of the most recognizable fish, especially among the non-scientific community, following the Disney movie “Finding Nemo” [2]. Even before the release of this film more than 15 years ago, the visual appeal and ability to complete their life cycle in captivity made clownfish a highly desired species in the marine aquarium trade [3,4]. For biologists, on the other hand, anemonefishes offer a unique opportunity to answer complex research questions about symbiosis, social dynamics, sex change, speciation, and phenotypic plasticity [1].

As for all anemonefishes, *A. ocellaris* is a protandrous hermaphrodite (i.e., male-to-female sex change) that lives in association with sea anemones [5–7]. It is a mutualistic relationship in which the host sea anemone provides food and shelter from predators [6,7], whilst the fish provides supplemental nutrition [8] and increased oxygen uptake by modulating water flow among the anemone’s tentacles [9,10]. The false clownfish is a generalist and can colonize up to four hosts: the magnificent sea anemone (*Heteractis magnifica*), the leathery sea anemone (*Heteractis crispa*), the giant carpet anemone (*Stichodactyla gigantea*), and Mertens’ carpet sea anemone (*Stichodactyla. mertensii*) [7]. Social groups typically consist of an adult breeding pair and several smaller (immature) juveniles ranked by size [11]. Within this social group a large dominant female is followed by a sub-dominant male, so that if the female is removed, the male changes sex and the largest non-breeder matures into a breeding male (sequential hermaphroditism) [12]. Within the Amphiprioninae subfamily, the two clownfish, *A. ocellaris* and *A. percula* form a separate clade that diverged early in the phylogeny, thus making them attractive for studies of the evolutionary history and adaptive radiation of anemonefishes [13,14]. Both species share behavioral and ecological traits and have similar body coloration patterns (i.e., bright orange with three vertical white bars) that set them apart from other anemonefishes. The major difference between these two species of clownfish is their differing geographic distributions, with *A. percula* restricted to the Great Barrier Reef, Papua New Guinea, and the Solomon Islands, whilst *A. ocellaris* can be found across the Indo-Malaysian region, from the Ryukyu Islands of Japan, throughout South East Asia, and south to north western Australia [7,13,15–18].

Until now, genome assemblies of at least ten anemonefish species including *A. ocellaris* have been published [19–22]. Yet, except for *A. percula*, these genomes are mainly based on Illumina short-read technology and are therefore highly fragmented, resulting in multiple gaps and mis-assemblies. However, third-generation sequencing platforms such as Pacific Biosciences, produce longer reads (5 to 60 kb) that enhance the continuous assembly of genome sequences [23,24]. This makes it possible to assemble complex regions of genomes, thus improving our ability to decipher genomic structures (such as chromosome rearrangements) and long-range regulatory analysis [25]. For example, 29 % of N-gaps in the human reference genome (GRCh38) could be filled with PacBio long reads [26]. In the case of *A. ocellaris*, the inclusion of Nanopore long reads together with Illumina data led to a 94 % decrease in the number of scaffolds, an 18-fold increase in scaffold N50 (401.72 kb), and a 16% improvement in genome completeness [22]. The PacBio long-read assembly of the *A.percula* genome further emphasized the power of long-read technology, with an initial contig assembly N50 of 1.86 Mb, further anchored into the chromosome-scale assembly (scaffold N50 of 38.4 Mb) [19].

Here, we constructed a high-quality chromosome-scale genome assembly and gene annotation for the false clownfish *A. ocellaris.* Using a combination of PacBio and Hi-C sequencing, we produced a *de novo* assembly comprised of 1,551 contigs with an N50 length of 863,854 bp that were successfully anchored into 24 chromosomes of 856,612,077 bp. We annotated 26,797 protein-coding gene models with the proportion of conserved actinopterygian genes reaching 96.62 %, making the quality and completeness of our genome better than previously published anemonefish genomes. Moreover, transcriptomic analysis detected 1,237 genes with absolute tissue-specific expression. A further comparative genomic approach identified genomic elements conserved only in the *A. ocellaris/A. percula* branch but not in other anemonefishes, many of which are associated with genes involved in nervous system functioning and were differentially expressed in brain tissues. Access to a high-quality chromosome-scale genome for *A. ocellaris* will not only unlock the potential to gain new insight into the evolutionary processes underlying the radiation of anemonefishes, but it will also facilitate our understanding of how the genomic architecture of this diverse group of reef fishes evolved. Ultimately, our work adds to the growing body of high-quality fish genomes critical to study genetic, ecological, evolutionary, and developmental aspects of marine fishes in general.

## METHODS

### Specimen collection and nucleic acid sequencing

Three adult *A. ocellaris* clownfish (one female and two males) were collected from 5 m depth in Motobu, Okinawa (26° 71’ 29.83” N, 127° 91’ 57.51” E) on March 25, 2020. Fish were kept under natural conditions at the OIST Marine Science Station in a 270 L (60 × 90 × 50 cm) tank until May 19, 2020. Individuals were euthanized following the guidelines for animal use issued by the Animal Resources Section of OIST Graduate University. Tissues for genome sequencing were snap frozen in liquid nitrogen and then stored at −80 □ until further processing. Genomic DNA was extracted from a male clownfish using a Qiagen tissue genomic DNA extraction kit (Hilden, Germany) and sequenced at Macrogen (Tokyo, Japan). For genome assembly, we sequenced genomic DNA from the brain tissue of the same male fish using two different platforms: PacBio Sequel II and Illumina NovaSeq6000 (Table S1). For long-read sequencing, 8 μg of genomic DNA was used to generate a 20 kb SMRTbell library according to the manufacturer’s instructions (Pacific Biosciences, CA, USA). Briefly, a 10 μL SMRTbell library was prepared using a SMRTbell Express Template Prep Kit 2.0 and the resulting templates were bound to DNA polymerases with a Sequel II Binding Kit 2.0 and Internal Control Kit 1.0. Sequencing on the PacBio Sequel II platform was performed using a Sequel II Sequencing Kit 2.0 and a SMRT cells 8M Tray. SMRT cells using 15 hr movies were captured. For short-read sequencing, a library was prepared from 1 μg of genomic DNA and a TruSeq DNA PCR-free Sample Preparation Kit (Illumina, CA, USA). Paired-end (151 bp per read) sequencing was conducted using a NovaSeq6000 platform (Illumina, CA, USA).

Hi-C reads were also sequenced to capture chromatin conformation for chromosome assembly. Liver (> 100 mg) tissue from another male fish was snap frozen and stored at −80 □ (Table S1). The tissue was sliced into small pieces using a razor blade (to increase the surface area for efficient cross-linking), resuspended in 15 mL of 1% formaldehyde solution, and then incubated at room temperature for 20 min with periodic mixing. Glycine powder was added to the solution for a final concentration of 125 mM followed by a 15 min incubation at room temperature with periodic mixing. Samples were spun down at 1,000 *g* for 1 minute, the supernatant was removed, and the tissue was rinsed with Milli-Q water. Tissues were then ground into a fine powder using a liquid nitrogen-chilled mortar and pestle. Powdered samples were collected and stored at −80 °C. Chromatin isolation, library preparation and Hi-C sequencing was performed by Phase Genomics (WA, USA). Following the manufacturer’s instructions, a Proximo Hi-C 2.0 Kit (Phase Genomics, WA, USA) was used to prepare the proximity ligation library and process it into an Illumina-compatible sequencing library. Hi-C reads were sequenced on an Illumina NovaSeq6000 platform to generate 150 bp paired-end reads.

Tissues for transcriptome sequencing were dissected from two individuals (one male and one female) and stored in RNAlater stabilization solution (Sigma Life Science, MO, USA) at −80 □. Transcriptome library preparation and sequencing were performed by Macrogen (Tokyo, Japan). Briefly, mRNA was extracted from brain optic lobe, caudal fin, eye, gill, gonads (from male and female fish), intestine, kidney, liver, the rest of the brain, skin (from orange and white bands), and stomach tissues using a Qiagen RNeasy Mini Kit (Hilden, Germany). Only high-quality RNA samples with an RNA integration number (RIN) > 7.0 were used for library construction. Libraries were prepared with 1 μg of total RNA for each sample using a TruSeq Stranded mRNA Sample Prep Kit (Illumina, CA, USA). Paired-end sequencing (151 bp) was conducted on a NovaSeq6000 machine.

### Chromosome-scale genome assembly of *A. ocellaris*

Prior to *de novo* assembly genome size was estimated using Jellyfish v2.3.0 [27] with *k*-mer = 17 and default parameters, and GenomeScope v1.0 [28] with default parameters. Quality-trimmed Illumina short reads obtained from Trimmomatic v0.39 [29] using the parameter set ‘ ILLUMINACLIP:TruSeq3-PE.fa:2:30:10:8:keepBothReads LEADING: 3 TRAILING: 3 MINLEN:36’ were used as input for Jellyfish. Genomic contigs were assembled using the FALCON software version as of September 28, 2020. For chromosome-scale assembly, initial contigs obtained from FALCON-phase were scaffolded with Phase Genomics’ Proximo algorithm based on Hi-C chromatin contact maps. In brief, the processed Hi-C sequencing reads were aligned to the Falcon assembly with BWA-MEM [30] using the −5 SP and -t 8 options. PCR duplicates were flagged with SAMBLASTER v0.1.26 [31] and subsequently removed from all following analyses. Non-primary and secondary alignments were filtered using SAMtools v1.10 [32] with the -F 2304 flag. FALCON-Phase [33] was then used to correct phase switching errors in the scaffolds obtained from FALCON-Unzip [34].

A genome-wide contact frequency matrix was built from the aligned Hi-C read pairs and normalized by the number of DPNII restriction sites (GATC) on the scaffolds, as previously described [35]. A total of 40,000 individual Proximo runs were performed to optimize chromosome construction. Juicebox v1.13.01 [36] was used to correct scaffolding errors and FALCON-Phase was again used to correct phase switching errors detectable at the chromosome level but not at the scaffold level. Local base accuracy in the long read-based draft assembly was improved with Illumina short reads using Pilon v1.23 [37]. Quality-trimmed Illumina short reads obtained from Trimmomatic v0.39 [29] using the same parameter set described above were aligned to the proximo-assembled chromosome-scale genome with Bowtie2 v2.4.1 [38] using the default settings. SAM files were converted to BAM files with SAMtools v1.10 [32] and then used as input for Pilon. Error correction was completed using five iterations of Pilon.

To calculate the overall mean genome-wide base level coverage, PacBio reads were aligned to the assembled chromosome sequences using Pbmm2 v1.4.0 (https://github.com/PacificBiosciences/pbmm2). Per-base coverage of aligned reads across entire chromosomal sequences was obtained using the BEDTools v2.30.0 [39] genomeCoverageBed function. Finally, we compared the quality of our genome assembly to three other published *A. ocellaris* genome sequences [21,22] using Quast v5.0.2 [40].

### Prediction of gene models in *A. ocellaris*

Repetitive elements in the *A. ocellaris* genome were identified *de novo* using RepeatModeler v2.0.1 [41] with the parameter -LTRStruct. RepeatMasker v4.1.1 [42] was then used to screen known repetitive elements with two separate inputs: the RepeatModeler output and the vertebrata library of Dfam v3.3 [43]. The two output files were validated, merged, and redundancy was removed using GenomeTools v1.6.1 [44].

BRAKER v2.1.6 [45] was then used to annotate candidate gene models of *A. ocellaris.* For mRNA evidence for gene annotation, transcriptomic reads (Table S1) were trimmed with Trimmomatic v0.39 [29] using the parameter set mentioned above and mapped to the chromosome sequences with HISAT2 v2.2.1 [46] using the ‘-dta’ option. SAM files were then converted to BAM format using SAMtools v1.10 [32]. For protein evidence, manually annotated and reviewed protein records from UniProtKB/Swiss-Prot (UniProt Consortium, 2021) as of January 11, 2021 (563,972 sequences) in addition to the proteomes of the false clownfish (*A. ocellaris:* 48,668), zebrafish (*Danio rerio:* 88,631), spiny chromis damselfish (*Acanthochromis polyacanthus:* 36,648), Nile tilapia (*Oreochromis niloticus:* 63,760), Japanese rice fish (*Oryzias latipes:* 47,623), rainbow fish (*Poecilia reticulata:* 45,692), bicolor damselfish (*Stegastes partitus:* 31,760), tiger puffer (*Takifugu rubripes:* 49,529), and Atlantic salmon (*Salmo salar:* 112,302) from the NCBI protein database

(https://www.ncbi.nlm.nih.gov/protein) were used. Only gene models with evidence support (mRNA or protein hints) or with homology to the Swiss-Prot protein database [47] or Pfam domains [48] identified by Diamond v2.0.9 [49] and InterProScan v5.48.83.0 [50], respectively, were added to the final gene models. Benchmarking Universal Single-Copy Orthologs (BUSCO) v4.1.4 [51] with the Actinopterygii-lineage dataset (actinopterygii_odb10) was used for quality assessment of gene annotation. Finally, for functional annotation of predicted gene models, NCBI BLAST v2.10.0 [52] was used with the NCBI non-redundant protein database (*nr*) as the target database. Gene Ontology (GO) terms were assigned to *A. ocellaris* genes using the ‘gene2go.gz’ and ‘gene2accession.gz’ files downloaded from the NCBI ftp site (https://ftp.ncbi.nlm.nih.gov/gene/DATA/) and the BLAST output.

### Assembly and annotation of the mitochondrial genome

The mitochondrial genome of *A. ocellaris* was assembled using Norgal v1.0.0 [53] with quality trimmed Illumina genomic reads. MitoAnnotator v3.67 [54] was used to annotate the organelle genes. Annotated genes in this study were compared to previously published *A. ocellaris* genes using BLASTn v2.10.0 [52] with e-value 10^-4^ as a threshold to predict homology. Only the longest isoform of each gene model was used for the homology search.

### Analysis of gene expression

Transcriptomic reads from each tissue were processed with Trimmomatic v0.39 [29] using the parameter set mentioned above and mapped to the genome using HISAT2 v2.2.1 [46]. SAM files were then converted to BAM files using SAMtools v1.10 [32]. Expression levels were quantified and TPM (transcripts per million) was normalized with StringTie v2.1.4 [55]. Tissue-specify index (τ) was calculated for each gene using the R package tispec v0.99 [56], with the relationship between τ and TPM expression values visualized on a two dimensional histogram with ggplot2 v3.3.5 [57]. TPM expression values per tissue were visualized in an Upset plot with the UpSetR v1.4.0 package [58].

### Gene orthology and phylogenetic analyses

To identify evolutionary relationships between *A. ocellaris* and other Amphiprioninae species, two species combinations were used: (1) a dataset that includes 11 anemonefish proteomes, i.e. our *A. ocellaris* proteome and ten other anemonefishes (*Amphiprion akallopisos*, *Amphiprion bicinctus*, *Amphiprion frenatus*, *Amphiprion melanopus*, *Amphiprion nigripes*, *Amphiprion percula*, *Amphiprion perideraion*, *Amphiprion polymnus*, *Amphiprion sebae*, and *Premnas biaculeatus*) [19–21], and *A. polyacanthus* as a single outgroup species, and (2) a dataset comprised of all 11 anemonefishes, *A. polyacanthus*, and five additional outgroup species across the teleost phylogenetic tree: zebrafish (*D. rerio*), bicolor damselfish (*S. partitus*), Asian seabass (*Lates calcarifer*), Nile tilapia (*O. niloticus*), and southern platyfish (*Xiphophorus maculatus).* The proteomes of outgroup species were obtained as previously described [19]. In all cases, only the longest isoform of each gene model was utilized.

Ortholog gene relationships between all taxa were investigated using OrthoFinder v2.5.2 [59]. Proteins were reciprocally blasted against each other, and clusters of orthologous genes (i.e., genes descended from a single gene in the last common ancestor) were defined using the default settings. Phylogenetic relationships of fish species were then assessed based on concatenated multi-alignments of one-to-one orthologs. In brief, sequences of single-copy orthologs present in all species were first aligned using MAFFT v7.130 [60] using the options ‘-localpair -maxiterate 1,000 -leavegappyregion’, then trimmed with trimAl v1.2 [61] using the ‘-gappyout’ flag, and finally concatenated with FASconCAT-G [62].

Phylogenetic trees were first constructed based on maximum-likelihood criteria using the two datasets described above. The MPI version of RAxML v8.2.9 (raxmlHPC-MPI-AVX) [63] was executed using a LG substitution matrix, heterogeneity model GAMMA, and 1,000 bootstrap inferences. Next, a subset of proteins for each species that has a complete match to the Actinopterygii-lineage (actinopterygii_odb10) identified by BUSCO v4.1.4 [51] were selected, concatenated, and used to construct new maximum-likelihood and Bayesian trees. Bayesian tree reconstructions were conducted under the CAT-GTR model as implemented in PhyloBayes MPI v1.8 [64]. Two independent chains were run for at least 5,000 cycles and sampled every 10 trees. The first 2,000 trees were removed as burn-in. Chain convergence was evaluated so that the maximum and average differences observed at the end of each run were < 0.01 in all cases. Trees were visualized and re-rooted using iTOL v6.4 [65]. Branch supports in the phylogenetic trees were evaluated with the standard bootstrap values from RAxML and PhyloBayes for maximum-likelihood and Bayesian trees, respectively. Site concordance factors (i.e., the proportion of alignment sites that support each branch) were also evaluated using IQ-TREE v2.1.3 [66].

### Interspecies synteny

Patterns of synteny (i.e., the degree to which genes remain on corresponding chromosomes) and collinearity (i.e., in corresponding order) across all anemonefish genomes were investigated using the MCScanX toolkit [67]. Briefly, an all-versus-all BLASTp search (using the parameters ‘-evalue 10^-10^ -max_target_seqs 5’) was first performed to identify gene pairs among species. Synteny blocks between two species were then calculated using the following parameters: ‘-k 50 -g −1 -s 10 -e 1e-05 -u 10,000 -m 25 -b 2’. This approach identified collinear blocks that had at least ten genes with an alignment significance <10^-5^ in a maximum range of 10,000 nucleotides between genes. Results were visualized using SynVisio [68]. The divergence time between two species were obtained from the TimeTree database [69].

### Identification of conserved genomic elements

For whole genome alignment analysis, genome sequences and gene annotations of the previously selected 11 anemonefish species and *A. polyacanthus* were used. Repeat elements were identified using RepeatModeler v2.0.1 [41] and RepeatMasker v4.1.1 [42] as described above and then soft-masked using BEDTools v2.30.0 [39]. Repeat-masked genome sequences and phylogenetic trees constructed with RAxML v8.2.9 [63] were used as input for whole genome alignment with Cactus multiple genome aligner v1.3.0 [70]. Resulting HAL databases were converted to MAF format using hal2maf v2.1 [71] with the *A. ocellaris* genome as a reference. MAFFILTER v1.3.1 [72] was then used to exclude repetitive regions and short alignments (< 100 bp). RPHAST v1.6.11 [73] was used to identify conserved genomic elements in the *A. ocellaris/A. percula* branch from the alignment. Adjusted p-values of significant conservation for genomic elements were computed using the Benjamini & Hochberg method with the p.adjust function implemented in the R package stats v4.1.0 [74]. Genomic elements with an adjusted p-value < 0.05 were considered as significantly conserved. Genes close to these genomic elements were identified with the closestBed function of BEDTools v2.30.0 [39]. Conserved elements were visualized using Circos v0.69-8 [75]. Expression values of genes close to conserved elements in the *A. ocellaris/A. percula* branch were visualized with a heatmap using the heatmap.2 function in gplots v3.1.1 [76].

## RESULTS AND DISCUSSION

### Chromosome-scale genome assembly of *A. ocellaris*

To construct high-quality chromosomes of *A. ocellaris*, we first generated 12,376,320 PacBio long-reads (average read length 10,239 bp) and 672,631,646 Illumina short reads (read length 151 bp) from brain tissue of an adult *A. ocellaris* individual (Table S1). Prior to the *de novo* draft genome assembly, we investigated the global properties of the genome with Illumina short reads using Jellyfish v2.3.0 [27] and GenomeScope v1.0 [28]. At *k*-mer = 17, the heterozygosity of *A. ocellaris* genome inferred from short reads was 0.26 % and the estimated haploid genome size was 805,385,376 bp. The repetitive and non-repetitive regions of the genome were estimated to be 343,219,574 bp (42.62 %) and 462,165,802 bp (57.38 %), respectively.

After the phased FALCON assembly with PacBio long reads [34], we obtained the primary (1,551 sequences, 861,420,186 bp, N50: 863,854bp) and alternate (8,604 sequences, 679,345,988 bp, N50: 116,448 bp) haplotigs. To build the chromosome-scale assembly, 145,019,677 Hi-C read pairs (150 bp) were generated from liver tissue (Table S1), and the Proximo™ scaffolding platform (Phase Genomics, WA, USA) was employed to orient *de novo* contigs into the chromosomes. This resulted in 353 sequences (865,612,980 bp) that consisted of 24 chromosome sequences (856,672,469 bp) and 329 short scaffolds that were not placed into chromosomes (8,940,511 bp). To improve the quality of the chromosome assembly, we performed iterative error-correction on the 24 chromosome sequences with Illumina short reads using Pilon v1.23 [37]. At the 5^th^ iterative run, 97.94 % of the reads were aligned to the 24 chromosome sequences. Finally, we obtained 24 chromosomes, with a length ranging from 21,987,767 bp to 43,941,765 bp, totaling 856,612,077 bp (Figure 1). Overall GC content of the *A. ocellaris* genome was 39.58 %. The mean base-level coverage of the assembled chromosomes was 103.89 ×. Completeness of the genome assembly was assessed with BUSCO v4.1.4 [51] using the Actinopterygii-lineage dataset (actinopterygii_odb10). The overall BUSCO score was 97.01 % (Complete and single-copy BUSCOs: 96.21 %; Complete and duplicated BUSCOs: 0.8 %; Fragmented BUSCOs: 0.52 %; Missing BUSCOs: 2.47 %) (Table 1).

**Figure 1.**
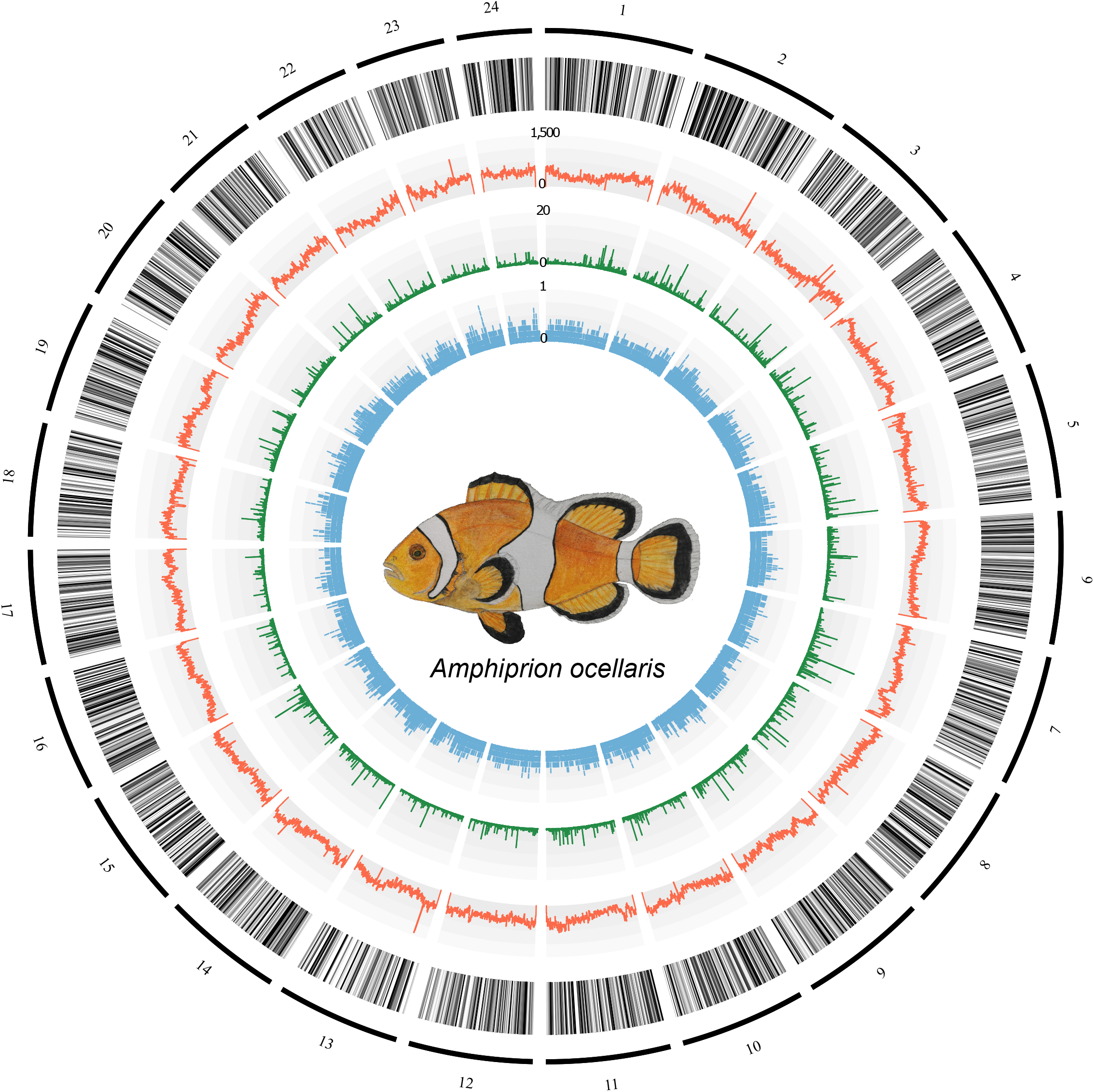
Chromosome architecture of the *Amphiprion ocellaris* genome. From the outermost layer inwards, each layer represents 1) chromosomes indicated by black lines and ordered by size, 2) genic regions (black colored), 3) the number of repeats per 100 kb (orange), 4) the PhyloP score calculated from the whole genome alignment of anemonefishes with *Acanthochromis polyacanthus* as an outgroup species (green), and 5) Tissue-specificity index (τ) of each gene (light blue).

**Table 1.** Statistics of the *Amphiprion ocellaris* chromosome-scale genome assembly and gene annotation.

|  |  |
| --- | --- |
| Chromosome assembly size | 24 sequences (856,612,077 bp) |
| Non-ATGC characters | 136,641 bp (0.02 %) |
| GC contents | 39.58 % |
| Mean base-level coverage | 103.89 × |
| Repeat contents | 44.7 % |
| BUSCO genome completeness | 3,531 (97.01 %) |
| Complete and single copy | 3,502 (96.21 %) |
| Complete and duplicated | 29 (0.8 %) |
| Fragmented | 19 (0.52 %) |
| Missing | 90 (2.47 %) |
| Number of protein coding genes | 26,797 |
| BUSCO gene annotation completeness | 3,517 (96.62 %) |
| Complete and single copy | 3,477 (95.52 %) |
| Complete and duplicated | 40 (1.1 %) |
| Fragmented | 38 (1.04 %) |
| Missing | 85 (2.34 %) |

Finally, we compared our chromosome-scale assembly to three other *A. ocellaris* draft genomes that have been previously published [21,22] (Table S2). In addition to large differences in size, which ranged from 744,831,443 to 880,704,246 bp, many misassembly events such as relocations, translocations, and inversions were observed in these other *A. ocellaris* genomes. This is likely due to the limitations of the short-read sequencing technologies upon which these assemblies were constructed.

### Prediction of *A. ocellaris* gene models

Repetitive elements in the *A. ocellaris* genome were examined by two approaches: (1) pattern matching using previously cataloged repetitive elements and (2) *de novo.* First the vertebrata repeat library from DFAM [43] was queried against the *A. ocellaris* genome sequences using RepeatMasker v4.1.1 [42]. We then identified 2,301 *de novo* repetitive elements using RepeatModeler v2.0.1 [41] and again searched for them in the *A. ocellaris* genome using RepeatMasker. A large fraction of the genome consisted of DNA transposons (24.11 %), long interspersed nuclear elements (LINEs, 7.67 %), long terminal repeats (LTRs, 3.77 %), and rolling-circle transposons (RCs, 1.55 %) (Figure 2 and Table S3). In total, 44.7 % (382,912,159 bp) of the whole genome was identified as repetitive elements. This is similar to the repeat content estimated from the unassembled short reads (42.62 %) using GenomeScope v1.0 [28] as described above. It should be noted though, that the sum of occupied percentages in the genome per repeat group is larger than the actual percentage (44.7 %) in the genome due to nested and overlapping repetitive elements (Figure 2).

**Figure 2.**
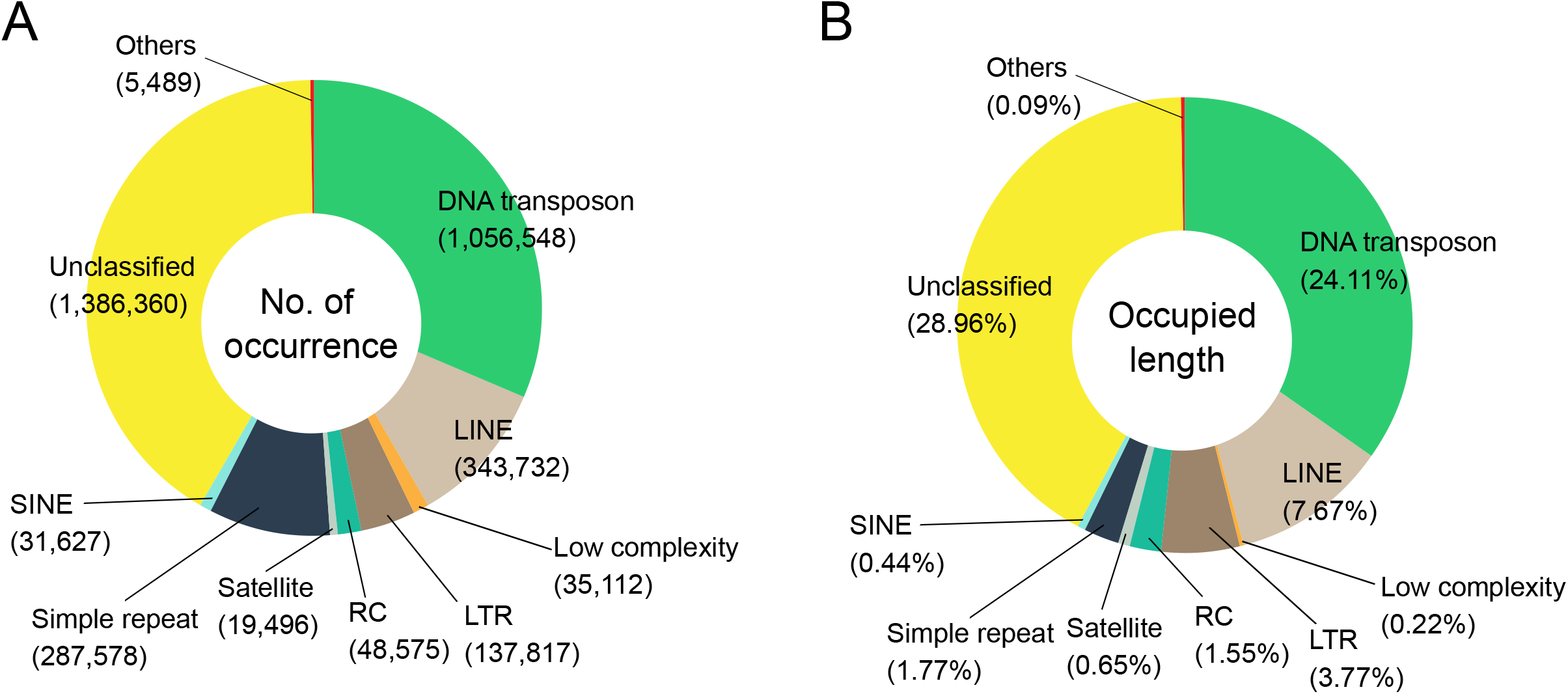
Repeat composition of the *Amphiprion ocellaris* genome. (A) The number of occurrences per repeat group classified by the DFAM database. (B) Occupied length in the genome per repeat group.

We then annotated the genome using BRAKER v2.1.6 [45] with mRNA and protein evidence. This evidence consisted of mapped transcriptomic reads sequenced from 13 tissues (Table S1), protein datasets from UniProtKB/Swiss-Prot [47], and the proteomes of 9 fish species. BRAKER predicted 26,433 gene models that were supported by either mRNA or protein hints, and 12,333 gene models with no evidence support. To account for the incompleteness of the evidence provided here and the gene annotation algorithm, we further added 364 nonsupport genes that have homology to the Swiss-Prot protein database and/or Pfam domains to the final gene models. This led to 26,797 final gene models from which 26,498 genes (98.88 %) had significant homology to the NCBI *nr* database (bit-score ≥ 50) and 21,230 genes (79.23 %) had at least one associated GO term. The completeness of our gene annotation was assessed using BUSCO v4.1.4 [51]. We obtained 96.62 % of completeness using the Actinopterygii-lineage dataset (Complete and single-copy BUSCOs: 95.52 %; Complete and duplicated BUSCOs: 1.1 %; Fragmented BUSCOs: 1.04 %; Missing BUSCOs: 2.34 %) (Table 1). This is higher than all other *A. ocellaris* and anemonefish gene annotations [19–22], in both the overall completeness and duplicated ratio, thus suggesting that the genome we present here is currently the best anemonefish genome annotation. Furthermore, our gene models include the majority (93.97 ~ 97.63 %) of gene models reported in previously published *A. ocellaris* genomes [21,22]. However, gene models from these studies include fewer gene models from this study (87.93 ~ 91.43 %), again indicating, that our gene models are the most comprehensive published to date (Table S4).

### Assembly and annotation of mitochondrial genome

We constructed the mitochondrial genome of *A. ocellaris* using Norgal v1.0.0 [53]. This resulted in a 16,649 bp circular mitogenome, which has the same length as another previously sequenced *A. ocellaris* mitochondrial genome (NCBI accession number: NC_009065.1). These two mitochondrial genomes showed 99.83 % sequence identity (16,621 of 16,649 bp) as calculated by BLASTn v2.10.0 [52]. MitoAnnotator v3.67 [54] was used to annotate the 37 organelle genes including 22 tRNA (Figure S1).

### Analysis of gene expression patterns across tissues

Gene expression levels of *A. ocellaris* genes were quantified using 13 tissue transcriptomes (Table S1). We investigated tissue-specificity of gene expression levels using the tau (τ) index as it is the most robust metric for identifying tissue-specific genes [77]. Given the range (0 ~ 1) of the τ index, we obtained 1,237 (4.62 %) absolutely specific genes (τ = 1; genes expressed only in one tissue), 5,302 (19.79 %) highly specific genes (0.85 ≤ τ < 1; genes highly expressed in a few tissues), and 3,431 (12.8 %) housekeeping genes (τ ≤ 0.2; genes expressed in nearly all tissues without biased expression) as defined by the R package tispec v0.99 [56]. Tissue-specificity of gene expression showed a negative correlation (Pearson’s correlation coefficient between τ and log_10_ maximum TPM value per gene = −0.46) with expression levels (Figure 3A), which is consistent with previous observations that highly tissue-specific genes tend to have lower expression levels [77,78].

**Figure 3.**
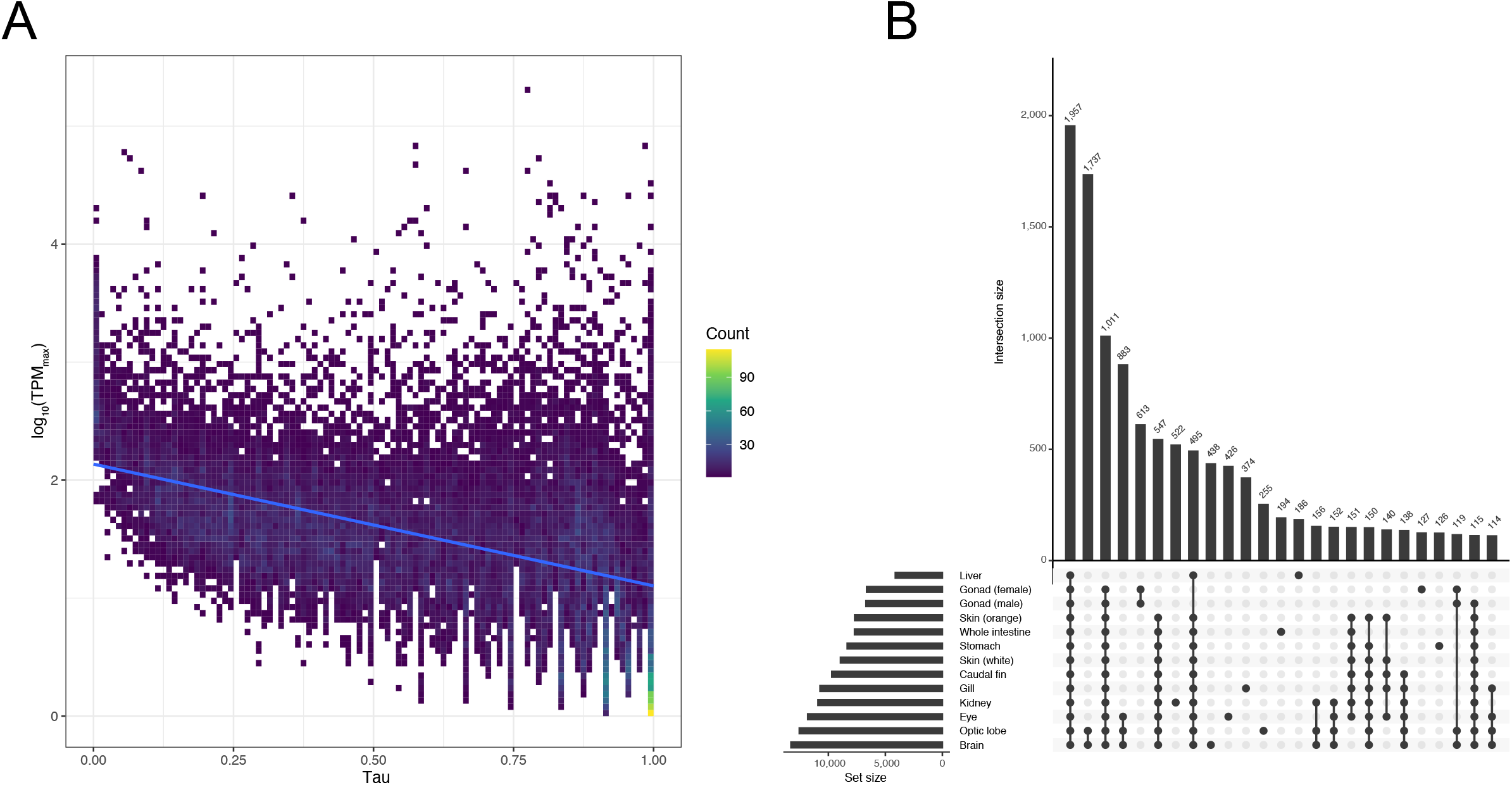
(A) Tissue-specificity of *Amphiprion ocellaris* gene expression. The maximum TPM (transcripts per million) values across tissues and tissue-specificity index (τ) was plotted in the two-dimensional histogram. Trendline (blue) was fit using the linear model. (B) Upset plot for the number of unique and shared genes expressed in different combinations of tissues. TPM values > 10 were used as the threshold for gene expression in the specific tissue. Intersection size represents the number of expressed genes in the designated sets.

Next, we checked gene expression patterns across tissues. After filtering for TPM ≥ 10, brain was the tissue with the highest number of expressed genes (*n* = 13,283) followed by optic lobe (*n* = 12,547) and eye (*n* = 11,809) (Figure 3B). This high number of genes expressed in the brain has also been reported in other vertebrates [78–80], and is most likely due to the complex role the brain has as the bodies control center. Furthermore, we observed that 1,957 genes were expressed in all 13 tissues sequenced here, and only 438 and 255 genes were exclusively expressed in the brain and optic lobe respectively (Figure 3B). Considering the high quality and similar numbers of transcriptomic reads per tissue generated in this study (Table S1), we are confident that these results represent the most accurate transcriptomic atlas for *A. ocellaris* to date.

### Phylogenetic analysis

Comparative analyses investigating the diversity and abundance of *A. ocellaris* gene families relative to other anemonefishes were performed using OrthoFinder v2.5.2 [59] with *A. polyacanthus* as an outgroup species (Table S5). Overall, most sequences (96.7%) could be assigned to one of 29,111 orthogroups, with the remainder identified as “unassigned genes” with no clear orthologs (Table S6). Fifty percent of all proteins were in orthogroups consisting of 12 or more genes (G_50_ = 12) and were contained in the largest 10,672 orthogroups. Further, 15,899 orthogroups were shared amongst all the species examined here, and from these, 12,765 consisted entirely of single-copy genes (Table S6). Interestingly, all trees (Figure 4 and Figure S2) obtained here using maximum likelihood or Bayesian inference approaches had the same topology and shared some similarities with aspects of earlier work [13,14,21]: a monophyletic *A. polymnus* and *A. sebae* group clustering with an Indian Ocean clade represented by *A. bicinctus* and *A. nigripes*, the skunk anemonefishes *A. akallopisos* and *A. perideraion*, an “ephippium complex” comprising *A. frenatus* and *A. melanopus*, and the monophyletic *A. ocellaris/A. percula* sister-species.

**Figure 4.**
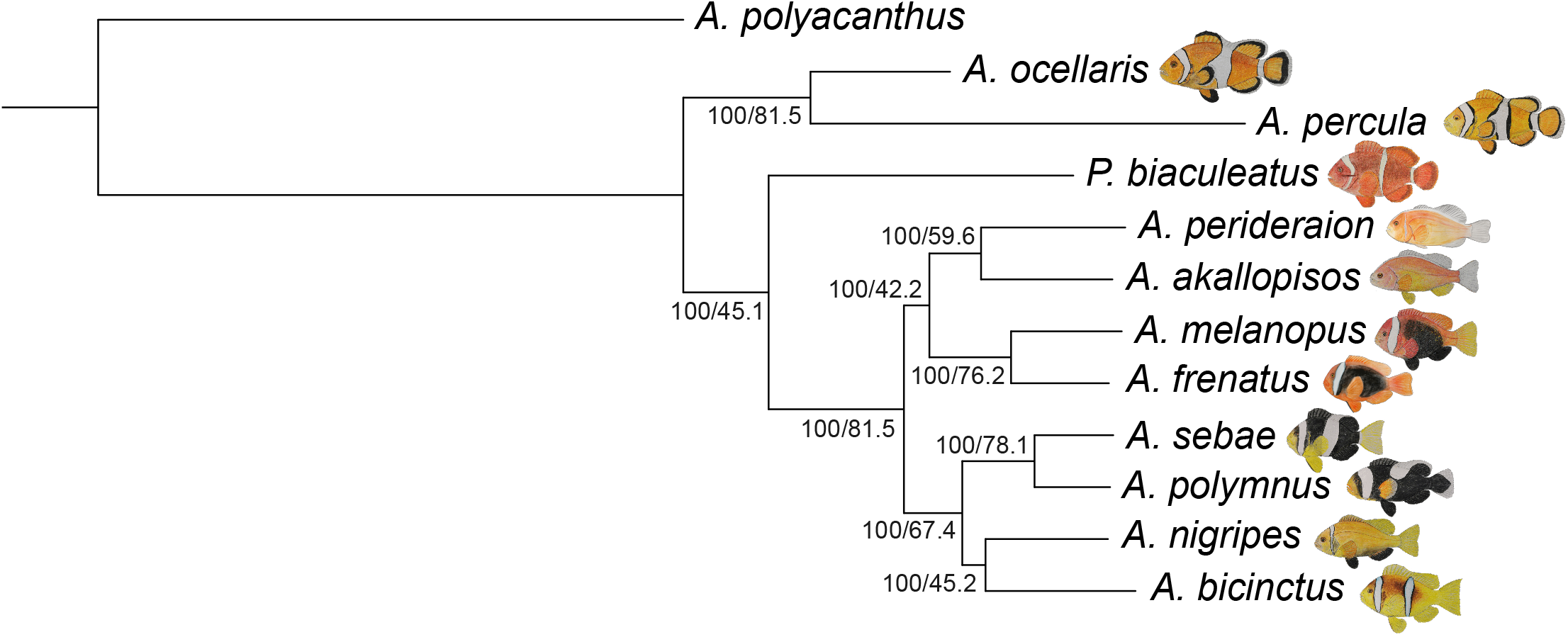
Phylogenetic reconstruction of the Amphiprioninae species tree using a maximum-likelihood approach. Numbers on each branching node are the bootstrap support (%) and the site concordance factor (%). These values were calculated only for non-outgroup species using the IQ-TREE algorithm.

Our tree diverges most dramatically from previous analyses in that *P. biaculeatus* was not positioned within the *A. ocellaris/A. percula* clade but became the root of all other anemonefishes with 100 % bootstrap support (Figure 4 and Figure S2). This topology has only been reported in two other studies that used mitochondrial genes to reconstruct anemonefish phylogenies [81,82]. Pomacentrids (and anemonefishes in particular) have long been a challenge in systematics due to their high diversity and intraspecific variation [83], therefore discordances in our tree may also stem from insufficient information (i.e., only 11 out of the 28 described species were used). Specifically, the inclusion of *Amphiprion latezonatus*, the sister group of all *Amphiprion* except for the *A. ocellaris/A. percula* clade, could be essential to resolve the molecular phylogeny of anemonefishes [13,14,82]. This topology could also be the result of gene choice, as incongruences between trees based on mitochondrial and nuclear data has previously been observed [14]. Yet, here, we used an alignment matrix consisting of more than twelve thousand single-copy genes (182,497 parsimony informative sites and 2.8% gaps) and still obtained this topology (Figure 4). This was further confirmed using BUSCO genes (Figure S2), predefined sets of reliable markers for phylogenetic inference [84].

We also observed weak support values using site concordance factors (i.e., the percentage of sites supporting a specific branch over 1,000 randomly sampled quartets) in some branches. For example, a support value of 45.89 % was recovered at the branching node of *P. biaculeatus* despite having 100 % bootstrap support (Figure 4 and Figure S2), thus suggesting high uncertainty. Still, despite the incongruence observed here, our phylogenetic reconstructions are based on large-scale genomic evidence, whilst other studies have used only a few genes. Although we are confident that our trees have a good resolution and represent one of the most enriched phylogenies for anemonefishes in terms of supporting genomic loci and reduced stochastic error, we are nonetheless cautious in our interpretation of the phylogenetic delimitation of species presented here. Certainly, establishing a well-resolved phylogeny of anemonefish, particularly the early divergent species (i.e., *P. biaculeatus*, the *A. ocellaris/A. percula* clade, and *A. latezonatus*), is critically important to understanding the evolution, genomic underpinning of their lifestyle (e.g., symbiosis with sea anemones, complex social structure) and fascinating biological features (e.g., pigmentation, sex change, aging).

### Whole genome synteny of anemonefishes

Syntenic blocks are often used to evaluate micro- and macro-scale patterns of evolutionary conservation and divergence among related species. Identifying conserved gene order at the chromosomal level among species furthers our understanding of the molecular processes that led to the evolution of chromosome structure across species [67,85]. Thus, here we used MCScanX [67] to investigate whole genome synteny among all species present. Overall, synteny patterns were consistent with the phylogenetic tree, in that closely related species had a higher number of conserved blocks than distant species (Table S7 and Figure S3), ultimately reflecting how gene gains or losses and sequence divergence increase proportionally with evolutionary time [85]. Yet, since all species studied here are still closely related, shared synteny among species pairs is considerably high. As expected, synteny between *A. ocellaris* and *A. percula* was much higher than comparisons to other anemonefishes (Table S7 and Figure S3). This analysis identified 175 syntenic blocks of 19,872 genes (Table S7) ranging from 11 to 1,010 gene pairs with 76.2 % of these being collinear (i.e., conserved order).

Although studying pairwise collinear relationships among chromosomal regions allows for the elucidation of gene family evolution, the alignment of multiple regions is even more important as it can reveal complex chromosomal duplication and/or rearrangement relationships [67]. Teleost fish genomes have been dynamically shaped by several forces (such as WGD and transposon activity). These in turn, led to various types of chromosome rearrangements either through differential loss of genes or formation of deletions, duplications, inversions and translocations, which together contribute to reproductive isolation and therefore might promote the formation of a new species [86]. Some chromosomal regions are translocated to new positions whereas others are inversed (Figure S4). While the information shown here is merely an initial overview of the large-scale synteny of the false clownfish and other anemonefishes, it is still an important first step in obtaining evolutionary insights into the Amphiprioninae lineage.

### Lineage-specific conserved genomic elements in the *A. ocellaris/A. percula* branch

Conserved genomic elements are relatively unchanged sequences across species. They are often parts of essential proteins or regulatory units and can be related to characteristics of specific lineages [86]. To identify the signature of such conserved elements in the *A. ocellaris* genome, we first attempted to identify conserved elements in *A. ocellaris* but not in other anemonefish using the PHAST program (see Methods) [73]. However, we were unable to identify genomic elements that were only conserved in this species, therefore we next sought to identify conserved elements shared by the two sister species of *A. ocellaris* and *A. percula*. We identified 91 conserved genome elements that showed significant conservation (adjusted p-value < 0.05).

To understand the possible role of these conserved elements, we investigated the function of 62 genes located around these elements (Table S8). It is interesting to note that at least 21 out of these 62 genes could be involved in neurological functions. For example, the *pcdh10* gene, encoding the protocadherin-10 protein, has been shown to be expressed in the olfactory and visual system of vertebrates as well as being involved in synapse and axon formation in the central nervous system [87]. Furthermore, a recent study also showed that mice lacking one copy of this gene have reduced social approach behavior [88]. Similarly, the *asic2* gene is expressed in the central and peripheral nervous system of vertebrates and its encoded protein, acid-sensing ion channel 2, is vital for chemo- and mechano-sensing the environment [89]. The neuronal pentraxin 2 protein, encoded by the *nptx2* gene, plays a role in the alteration of cellular activities for long-term neuroplasticity [90]. The *tafa5* gene encodes a neurokine involved in behavior related to spatial memory in mice [91] and peripheral nociception in zebrafish [92].

Additionally, these 62 genes also showed distinct expression patterns in *A. ocellaris* brain tissues (optic lobe and the rest of the brain) (Figure 5). Mean TPM expression level of these genes were 29.8 and 29.71 for the optic lobe and the other part of brain respectively, whereas mean TPM for other tissues was 14.17, potentially indicating different, yet unknown roles of these genes in the brain. Although further investigation is required, this data suggests that neuronal genes located around specifically conserved elements in the *A. ocellaris/A. percula* branch could represent genomic signatures related to the distinct ecology of these two sister species. Certainly, anemonefish societies are highly species-specific [17,93]. For example, while *A. percula* have a reduced range of movement, spending more time inside of their anemones and thus have a lower probability of social rank being usurped by outsiders, other species like *Amphiprion clarkii* have more opportunity for movement (due to their higher swimming abilities), increasing the probability of being taken over by outsiders so that the dominant individuals must display constant aggression to maintain control of their territory [94–96]. Future research should endeavor to better characterize these differences and investigate whether they are linked to the genes we have identified here.

**Figure 5.**
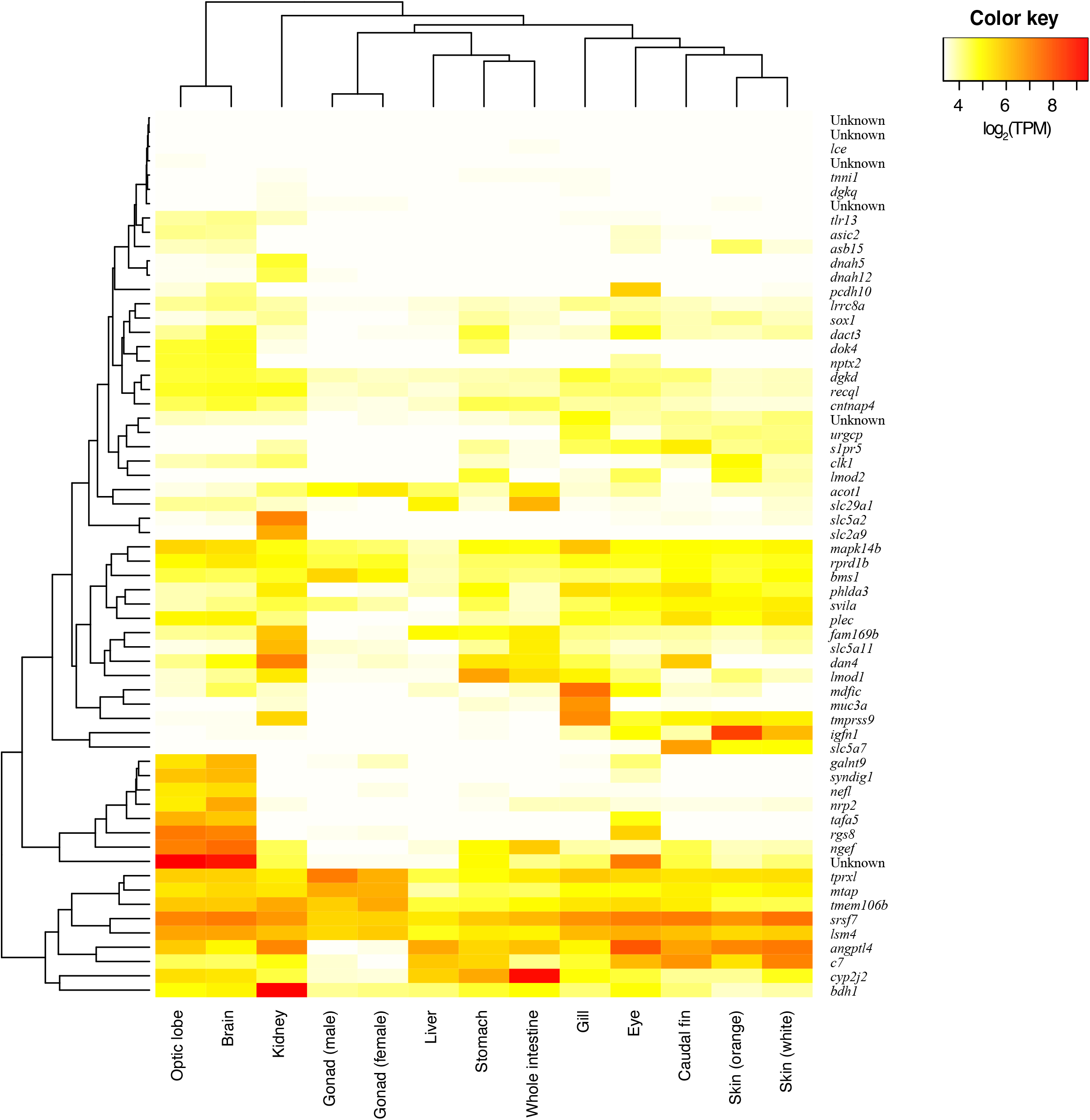
Gene expression levels of the 62 *Amphiprion ocellaris* genes nearest to the conserved elements in the *Amphiprion ocellaris/Amphiprion percula* branch across 13 tissues are shown. Color key indicates log_2_-transformed TPM values.

## CONCLUSIONS

Here, we assembled the highly contiguous and complete chromosome-scale genome of the false clownfish *A. ocellaris* by *de novo* assembly using PacBio long reads and Hi□C chromatin conformation capture technologies. We annotated 26,797 protein-coding genes with 96.62 % completeness of conserved actinopterygian genes, the highest level among anemonefish genomes available so far. We also identified tissue-specific gene expression patterns in *A. ocellaris*. Finally, we identified genomic elements conserved only in *A. ocellaris/A. percula*, which might underpin lineage-specific characteristics of these two species when compared to other anemonefishes. The high quality of our genome and annotation will not only serve as a resource to better understand the genomic architecture of anemonefishes, but it will further strengthen the false clownfish as an emerging model organism for molecular, ecological, developmental, and environmental studies of reef fishes.

## DATA AVAILABILITY STATEMENT

The sequencing reads generated in this study have been deposited in GenBank database under the BioProject ID PRJNA787397. The genome assembly and annotation are available in Dryad repository (https://datadryad.org/stash/share/UpzvIVKZOi21CcO38uwnPMgF1_ONpMM_LqX_S_0pS_oE).

## AUTHORS’ CONTRIBUTIONS

TR1 (Timothy Ravasi) and VL conceived the genome sequencing project. TR2 (Taewoo Ryu) designed the overall analysis. EK collected fish samples. BM and MI dissected tissues for sequencing. TR2 performed the overall computational analysis. MH performed to the orthology, phylogenetic, and synteny analyses. TR2 and MH wrote the manuscript with help from all authors. All authors read and approved the final manuscript.

## ACKNOWLEDGEMENTS

We thank Mr. Hidenori Kinjo for collecting the *A. ocellaris* samples used in this study and Dr. Konstantin Khalturin (OIST) for his help with the phylogenetic analysis. We also thank Mr. Lilian Carlu for the anemonefish drawings shown in Figures 1 and 4.

