## Supplementary Materials for "A chromosome-scale genome assembly of the false clownfish, *Amphiprion ocellaris*"

Table S1. Statistics for the sequencing dataset generated in this study.

| Data type | Sequencer | Fish ID | Tissue | Number of reads | Average read length (bp) | Size of dataset (bp) | BioSample ID |
| --- | --- | --- | --- | --- | --- | --- | --- |
| Genome | Pacbio Sequel II | 3 | Brain | 12,376,320 | 10,239 | 126,724,209,204 | SAMN23798798 |
| Genome | Illumina NovaSeq6000 | 3 | Brain | 672,631,646 | 151 | 101,567,378,546 | SAMN23798799 |
| Genome (Hi-C) | Illumina NovaSeq6000 | 2 | Liver | 290,039,354 | 150 | 43,505,903,100 | SAMN23798800 |
| Transcriptome | Illumina NovaSeq6000 | 1 | Brain | 45,640,648 | 151 | 6,891,737,848 | SAMN23798801 |
| Transcriptome | Illumina NovaSeq6000 | 1 | Caudal fin | 40,030,142 | 151 | 6,044,551,442 | SAMN23798802 |
| Transcriptome | Illumina NovaSeq6000 | 1 | Eye | 45,516,206 | 151 | 6,872,947,106 | SAMN23798803 |
| Transcriptome | Illumina NovaSeq6000 | 1 | Gill | 42,627,462 | 151 | 6,436,746,762 | SAMN23798804 |
| Transcriptome | Illumina NovaSeq6000 | 1 | Female gonad | 46,420,922 | 151 | 7,009,559,222 | SAMN23798805 |
| Transcriptome | Illumina NovaSeq6000 | 2 | Male gonad | 47,361,990 | 151 | 7,151,660,490 | SAMN23798806 |
| Transcriptome | Illumina NovaSeq6000 | 1 | Intestine | 47,565,906 | 151 | 7,182,451,806 | SAMN23798807 |
| Transcriptome | Illumina NovaSeq6000 | 1 | Kidney | 44,947,336 | 151 | 6,787,047,736 | SAMN23798808 |
| Transcriptome | Illumina NovaSeq6000 | 1 | Liver | 44,073,840 | 151 | 6,655,149,840 | SAMN23798809 |
| Transcriptome | Illumina NovaSeq6000 | 1 | Optic lobe | 47,344,796 | 151 | 7,149,064,196 | SAMN23798810 |
| Transcriptome | Illumina NovaSeq6000 | 2 | Orange skin | 50,188,374 | 151 | 7,578,444,474 | SAMN23798811 |
| Transcriptome | Illumina NovaSeq6000 | 2 | White skin | 40,142,226 | 151 | 6,061,476,126 | SAMN23798812 |
| Transcriptome | Illumina NovaSeq6000 | 1 | Stomach | 50,672,328 | 151 | 7,651,521,528 | SAMN23798813 |

Table S2. Comparison of the chromosome-scale assembly from this study to the previously reported genome assemblies for *Amphiprion ocellaris.*

|  | Previously published *A. ocellaris* genome assemblies | | |
| --- | --- | --- | --- |
|  | Tan et al. | Marcionetti et al. reference-guided assembly | Marcionetti et al. *de novo* assembly |
| Assembly statistics |  |  |  |
| Total length (>= 0 bp) | 880,704,246 | 798,423,830 | 744,831,443 |
| Total length (>= 50 kb) | 826,960,231 | 720,593,598 | 575,489,562 |
| # of scaffolds | 6,404 | 16,543 | 27,951 |
| Largest scaffold length | 3,111,502 | 1,729,050 | 1,238,133 |
| N50 | 401,715 | 246,482 | 136,417 |
| Misassembly statistics^*^ |  |  |  |
| # of relocation events | 5,986 | 20,991 | 4,215 |
| # of translocation events | 8,026 | 18,621 | 1,088 |
| # of inversion events | 107 | 2,459 | 28 |
| # of misassembled scaffolds^$^ | 4,123 | 8,026 | 3,360 |
| # of indels | 912,765 | 2,654,843 | 1,001,836 |
| # of fully unaligned scaffolds | 12 | 207 | 285 |
| Fully unaligned scaffold length | 56,591 | 261,822 | 661,296 |
| # of partially unaligned scaffolds | 2,751 | 5,671 | 5,380 |
| Partially unaligned scaffold length | 14,258,058 | 40,446,543 | 16,110,947 |
| Unaligned length | 14,314,649 | 40,708,365 | 16,772,243 |
| Total aligned length^*^ | 861,902,086 | 725,567,696 | 694,092,481 |

^*^ Alignment statistics were calculated against *A. ocellaris* genome assembly from this study.

^$^ The number of scaffolds with relocation, translocation, or inversion events

Table S4. Comparison of gene models from this study to the previously reported gene models for *Amphiprion ocellaris*

| Query gene models | Target gene models | # of query genes in target | # of target genes in query |
| --- | --- | --- | --- |
| This study | Marcionetti et al. *de novo* assembly | 23,562 (87.93 %) | 25,131 (96.85 %) |
| This study | Marcionetti et al. reference-guided assembly | 23,962 (89.42 %) | 28,109 (93.97 %) |
| This study | Tan et al. | 24,500 (91.43 %) | 26,594 (97.63 %) |
| Marcionetti et al. *de novo* assembly | Marcionetti et al. reference-guided assembly | 25,788 (99.38 %) | 28,959 (96.81 %) |
| Marcionetti et al. *de novo* assembly | Tan et al. | 25,180 (97.04 %) | 26,173 (96.08 %) |
| Marcionetti et al. reference-guided assembly | Tan et al. | 28,482 (95.22 %) | 26,464 (97.15 %) |

Only longest isoform per gene models were used for BLASTn search. Sequences with significant hit ( e-value < 10^-4^) were counted and percentage of genes in the entire gene set in each study were shown in the bracket.

Table S5. The number of gene models of each species used for phylogenomic analysis.

| Species | No. sequences |
| --- | --- |
| *A. polyacanthus* | 25,468 |
| *A. akallopisos* | 28,730 |
| *A. bicinctus* | 28,891 |
| *A. frenatus* | 26,917 |
| *A. melanopus* | 29,408 |
| *A. nigripes* | 28,558 |
| *A. ocellaris* | 26,797 |
| *A. percula* | 23,872 |
| *A. perideraion* | 29,014 |
| *A. polymnus* | 28,640 |
| *A. sebae* | 28,727 |
| *P. biaculeatus* | 28,170 |

Table S6. Summary of orthogroups identified by OrthoFinder.

| Number of species | 12 |
| --- | --- |
| Number of genes | 333,192 |
| Number of genes in orthogroups | 322,133 |
| Number of unassigned genes | 11,059 |
| Percentage of genes in orthogroups | 96.7 |
| Percentage of unassigned genes | 3.3 |
| Number of orthogroups | 29,111 |
| Number of species-specific orthogroups | 341 |
| Number of genes in species-specific orthogroups | 1,354 |
| Percentage of genes in species-specific orthogroups | 0.4 |
| Mean orthogroup size | 11.1 |
| Median orthogroup size | 12 |
| G50 (assigned genes) | 12 |
| G50 (all genes) | 12 |
| O50 (assigned genes) | 10,672 |
| O50 (all genes) | 11,133 |
| Number of orthogroups with all species present | 15,899 |
| Number of single-copy orthogroups | 12,765 |

Table S7. The number of shared syntenic blocks and ortholog gene pairs among 11 anemonefish species. Collinear synteny blocks were defined as a minimum of ten consecutive genes in both species. The numbers inside the bracket are the number of orthologous gene pairs within the synteny blocks between two species. Syntenic regions were not identified between any two pairs of *Amphiprion percula*, *Amphiprion perideraion*, nor *Amphiprion polymnus*.

|  | *A. akallopisos* | *A. bicinctus* | *A. frenatus* | *A. melanopus* | *A. nigripes* | *A. ocellaris* | *A. percula* | *A. perideraion* | *A. polymnus* | *A. sebae* | *P. biaculeatus* |
| --- | --- | --- | --- | --- | --- | --- | --- | --- | --- | --- | --- |
| *A. akallopisos* |  | 602  (9,247) | 507  (7,681) | 597 (9,184) | 594 (9,106) | 501 (7,460) | 432 (6,338) | 615 (9,414) | 595 (9,148) | 599 (9,183) | 583 (8,902) |
| *A. bicinctus* |  |  | 512  (7,695) | 586 (9,039) | 603 (9,295) | 493 (7,384) | 432 (6,341) | 596 (9,171) | 598 (9,213) | 595 (9,177) | 577 (8,820) |
| *A. frenatus* |  |  |  | 529 (7,994) | 505 (7,619) | 441 (6,532) | 395 (5,743) | 511 (7,733) | 504 (7,644) | 507 (7,661) | 504 (7,578) |
| *A. melanopus* |  |  |  |  | 587 (9,047) | 491 (7,353) | 425 (6,271) | 593 (9,162) | 599 (9,222) | 604 (9,270) | 573 (8,880) |
| *A. nigripes* |  |  |  |  |  | 492 (7,344) | 433 (6,327) | 590 (9,056) | 602 (9,234) | 592 (9,116) | 571 (8,736) |
| *A. ocellaris* |  |  |  |  |  |  | 175 (19,872) | 490 (7,349) | 503 (7,482) | 499 (7,453) | 484 (7,255) |
| *A. percula* |  |  |  |  |  |  |  | 0 (0) | 0 (0) | 426 (6,283) | 427 (6,279) |
| *A. perideraion* |  |  |  |  |  |  |  |  | 0 (0) | 590 (9,095) | 581 (8,878) |
| *A. polymnus* |  |  |  |  |  |  |  |  |  | 630 (9,652) | 571 (8,878) |
| *A. sebae* |  |  |  |  |  |  |  |  |  |  | 570 (8,736) |

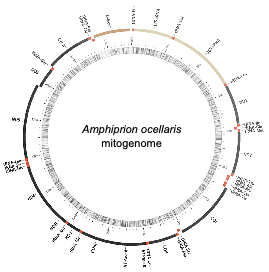

Figure S1. Mitochondrial genome annotation of *Amphiprion ocellaris*. The inner circle represents GC % per 5 bp of the mitogenome.

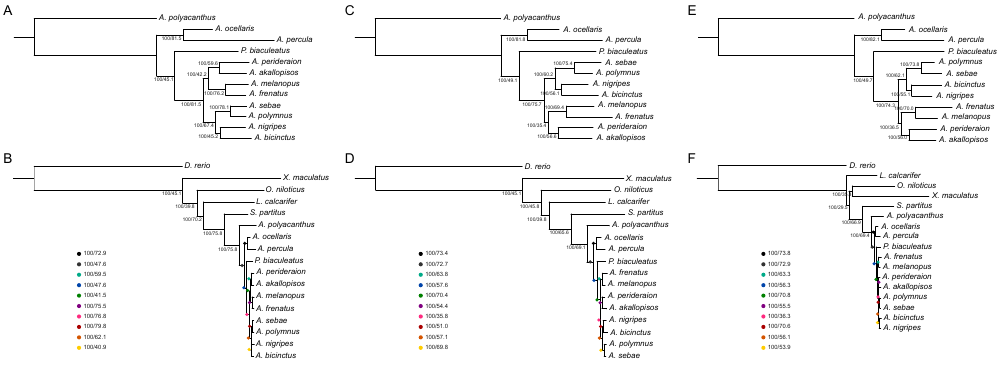

Figure S2. Phylogenetic trees constructed using (A) RAxML with 12,765 single copy genes from all 11 anemonefishes and their closest outgroup species *Acanthochromis polyacanthus*, (B) RAxML with 3,543 single copy genes from all 11 anemonefishes and six other fish species from across the teleost phylogenetic tree, (C) RAxML with 2,292 BUSCO genes from the same group of species used in A, (D) RAxML with 996 BUSCO genes from the same group of species used in B, (E) PhyloBayes with BUSCO genes from the same group of species used in C, and (F) PhyloBayes with BUSCO genes from the same group of species used in F. Numbers on each branching point are the bootstrap support (%) and the site concordance factor (%). These values were calculated only for non-outgroup species using the IQ-TREE algorithm.

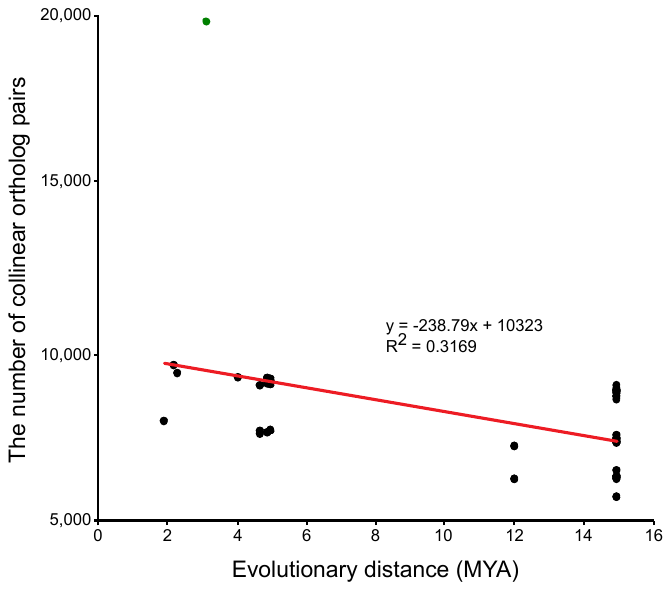

Figure S3. Negative correlation between the divergence time and the number of synteny blocks between species pairs. The species pairing between *Amphiprion ocellaris* and *Amphiprion percula* is marked in green.

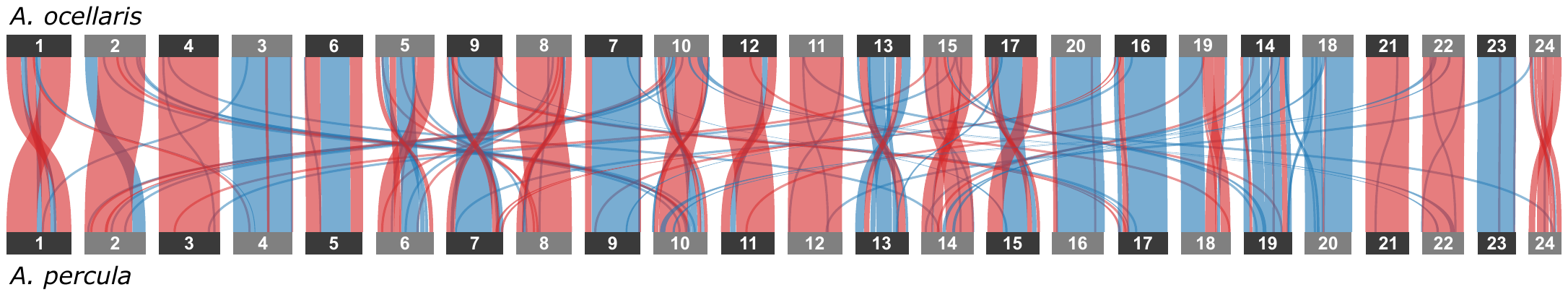

Figure S4. Dual synteny plot between all 24 chromosomes from *Amphiprion ocellaris* and *Amphiprion percula*. Chromosomal rearrangements such as translocations and inversions are shown as red ribbons, whereas blue ribbons represent unchanged regions.
