## Supplementary figures and images for "A chromosome-scale genome assembly of the false clownfish, *Amphiprion ocellaris*"

### Figure S2

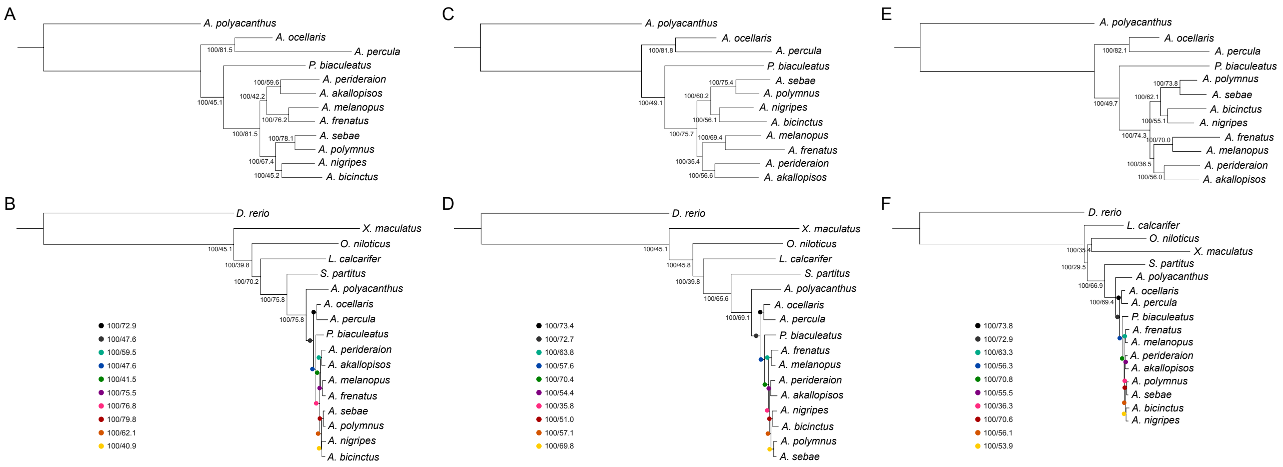

### Figure S3

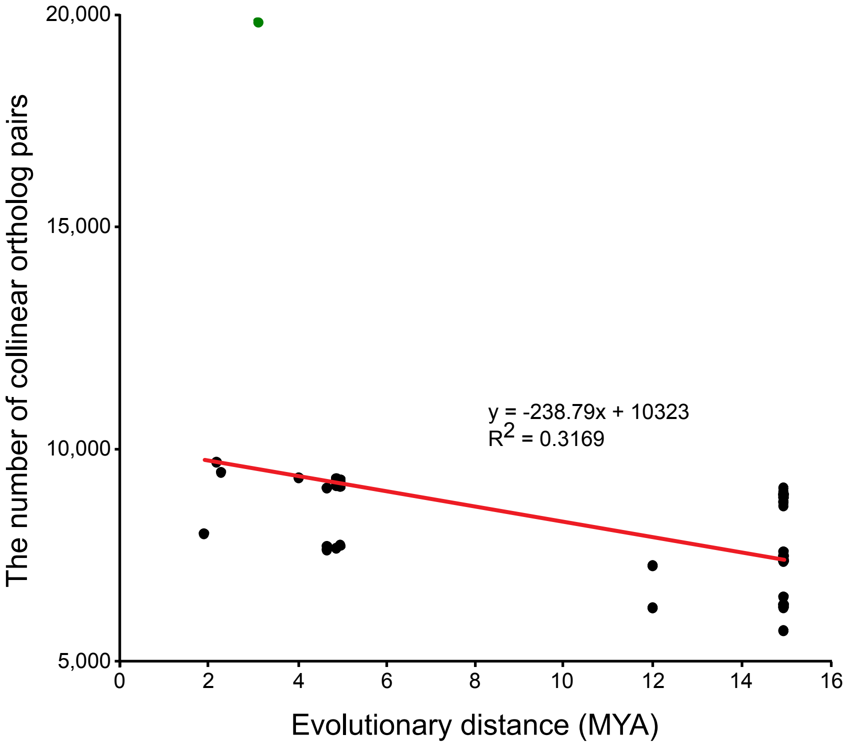

### Figure S4

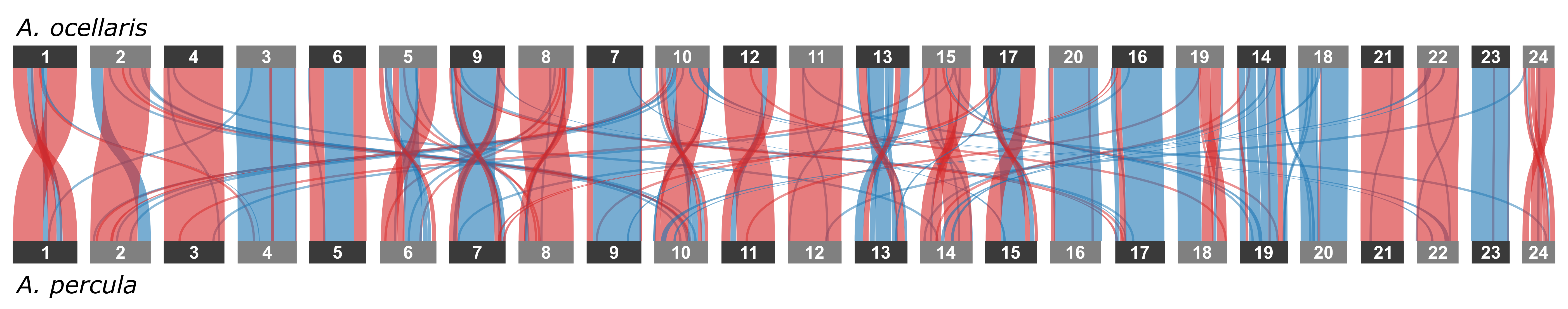
